# Furfural residues as soil amendments for long-term seed corn production: effects on community composition and diversity of rhizosphere microbiota

**DOI:** 10.1101/2020.11.25.397133

**Authors:** Yunchen Zhao, Guangquan Chen, Yuru Chen, Xuelin Song, Zhanwen Xiao, Zhijiang Wang, Wenjiang Fu

**Author notes:** **Corresponding author:** Yunchen Zhao, **Email address:**.

## Abstract

Furfural residues’ s influence on maize rhizosphere microbiota subjected to long-term monocropping is poorly understood. In this study, high-throughput sequencing was employed to investigate the rhizosphere microbiota composition and its variation under long-term field monocropping for maize seed production. The results showed that unplanted, chemical fertilizer (only) treated soil, and furfural residue treated soil as well as seasonal soil batches recruited distinct rhizosphere microbiota. Microbial community diversity increased, and many operational taxonomic units (OTUs) were induced in the rhizosphere soil. Maize plants grown under field conditions were preferentially colonized by Ascomycota and Zygomycota in the unplanted soil, Ascomycota and Mortierellomycota in chemical fertilizer-treated soil, and Ascomycota and Basidiomycota in frufural residue-treated soil. Some potential pathogens, such as *Alternaria, Trichocladium, Bipolaris, Solicoccozyma* and *Cladosporium* were not detected, while beneficial microbes, such as *Penicillium, Schizothecium* and *Rhizophlyctis* were present. Acidobacteria and Bacteroidetes increased, Actinobacteria and Firmicutes decreased in furfural residue treated soil. The core bacteria detected after long-term cropping were *MND1, RB41, UTCFX1, Nitrospira, Cellvibrio, Adhaeribacter* etc. The relative abundances of *Clostridium, Pseudarthrobacter* and *Roseiflexus* decreased; *Haliangium, Nitrospira* and *MND1* increased; *Pirellula, Ellin6067* and *Luteimonas* reduced in different seasonal soil batches. Amendment with furfural residue promoted the development of beneficial microbes and decreased the abundance of pathogens after different continuous cropping years. The amendment increased cellulose-degrading and complex carbon-decomposing microbes, decreased the number of reductive substance-decomposing microbes, which led to microbial community structure shift over time. Amendment with furfural residue improved the rhizosphere environment, which will in turn improve plant growth. Furfural residue can be used as a soil amendment to control soil-borne diseases and to establish beneficial soil microbes.

**Importance:** Continuous monoculture of maize seed production led to reduction in plant nutrient absorption, destruction of soil structure, high incidence of soil-borne plant diseases, and decrease in crop yield. Traditional organic fertilizers are either unavailable or unbalanced for intensive cultivation. Furfural residues are acidic and carbon-rich making it a promising organic alternative to chemical fertilizers to improve seed corn production. Amendment with furfural residue may promote the microbiota in rhizosphere soil. In addition, the amendment may increase cellulose-degrading and complex carbon-decomposing microbes, which led to a shift in microbial community structure over time. Amendment with furfural residue may improve the rhizosphere environment, which will in turn improve plant growth, control soil-borne diseases and establish beneficial soil microbes.

## 1. Introduction

Continuous monoculture of maize for seed production prevails in Northwest China. This farming has deleterious effects, including reduction in plant nutrient absorption (1), destruction of soil structure, high incidence of soil-borne plant diseases, and decrease in crop yield (2). Organic amendments improve structure and nutrient status of soil, supply food resources for microorganisms (3,4), and enhance microbial populations in rhizosphere (5,6). However, traditional organic fertilizers are either unavailable or unbalanced. Moreover, improper manure management introduces zoonotic pathogens (7) or spread antibiotic resistance genes (8,9), that in turn cause diverse environmental problems (10,11). Therefore, new organic sources have drawn substantial interest.

Hexi Corridor, particularly Zhangye City, with adequate solar and heat resources and broad area of flat lands, is an important corn seed production base. These region contributes to two-thirds of the total national seed supply. Large quantities of residues are generated from furfural production industries. These residues have been accumulated that lead to different environmental issues, such as pollution, in Northwest China.These residues are acidic (pH 2–3) and carbon-rich (cellulose and lignin) (12) making it a promising organic alternative to chemical fertilizers to improve seed corn production (1). However, the effects of furfural residue as soil amendments on the community diversity and structure of soil microbes remain largely unexplored. No studies have so far investigated the effects of furfural residues on rhizosphere microbes during long-term continuous cropping of maize for seed production. This study aims to (1) characterize the microbial composition in maize rhizosphere at different growth stages after the application of furfural residue; (2) reveal the population dynamics, diversity and structure of microbes in the rhizosphere during long-term continuous cropping after the application of furfural residue; and (3) explore the relationship between furfural residue and pathogens including soil-borne pathogens in the rhizosphere.

## 2. Materials and methods

### 2.1. Site description

The study was conducted at Dabai Village of Zhangye (Gansu Province, China) where more than 10 companies with nearly 1300 hectares of fields operated to produce corn seeds on an alkali irrigated desert soil. The properties of the soils are provided in Supplementary Table S1). The annual temperature, evaporation, and precipitation were approximately 4.2°C, 2100 mm, and 220 mm, respectively. Six fields with a planting history of 5 years (A), 10 years (B), 15 years (C), 20 years (D), 25 years (E) and 30 years (F) were selected for the study. The geographic location of these fields is provided in Supplementary Table S2.

### 2.2. Field experiments

All the fields were split in half; one was treated with furfural residue (k) and the other with chemical fertilizer only (c). All the fields were managed identically, and all the plots were irrigated separately. Furfural residue was neutralized with lime powder at the rate of 1.5 kg lime/100 kg residue and was applied at the rate of 22.2 t ha^-1^ one month before planting. The properties of the furfural residue after neutralization are shown in Supplementary Table S3. Furfural residue was plowed into the 10–20 cm soil layer, and the surface was raked until smooth to maintain moisture. Chemical fertilizers (420 kg ha^-1^ N and 380 kg ha^-1^ P_2_O_5_) were applied to all the fields. Urea (N, 46%) and phosphate diammonium (N, 18% and P_2_O_5_, 46%) were used as the fertilizers. The entire dose of P_2_O_5_ and one-third of N were applied as base fertilizers before planting, and one-third each of N fertilizer was applied as topdressing at bell mouth and milk-ripe stages. Mulch (70 cm width) was applied one week after irrigation, and parents of the maize cultivar Zhengdan958 (Zheng58/Chang7-2) were planted. Female parents were planted on April 20, and male parents at two different stage were planted 10 days and 20 days later in a flashing star and a triangular-symmetric fashion, respectively, to ensure synchronized flowering.

### 2.3. Soil sample collection

Soil samples from fields at different years of continuous cropping were collected before planting and fertilization on March 15 (unplanted soil). Each sample comprised a mixture of ten samples (0–20 cm soil layer). Rhizosphere samples were collected after planting and fertilization at bell mouth stage (July 18) from fields at different years of continuous cropping (rhizosphere soil) after removing the adhering soil by vigorously shaking the maize roots. The root hairs, stones and plastic pieces were carefully removed by hand. Each sample comprised rhizosphere soil from 10 plants. These samples (approximately 5 g each) were placed on ice and transferred to the laboratory within 1 hour for further testing. Triplicates were maintained per sample. Rhizosphere soil samples were collected at seedling stage (May 10) from fields treated with furfural residue at seedling as a seasonal batch for further analysis.

### 2.4. DNA extraction, PCR amplification and illumina Miseq sequencing of the soil

The DNA of samples was extracted by using the DNA Isolation Kit and the genomic DNA was amplified by PCR using the 16S r RNA gene V4V5 region primers (515 a – 806 a) and ITS1/ITS2(V4 – V5). The primers sequence was 515F: GTGCCAGCMGCCGCGGTAA, 907R: CCGTCAATTCMTTTRAGTTT, ITS3F: 5′-GCATCGATGAAGAACGCAGC-3′ and ITS4R: 5′-TCCTCCGCTTATTGATATGC-3′. PCR conditions: initialization for 5 min at 94°C, followed by 30 cycles denaturation at 94 °Cfor 30s, 30 s annealing at 52°C, and extension at 72°C for 30 s.The PCR instrument was BioRad S1000 (Bio-Rad Laboratory, CA, USA). All reactions ended with a final 72°C extension for10 min. The amplified product was electrophoresed in 1.5% of agarose gels at 100V for 30 min. PCR products were mixed in equidensity ratios according to the GeneTools Analysis Software. Then, the mixture of PCR products was purified by the EZNA Gel Extraction Kit (Omega, USA). Sequencing libraries were generated using NEBNext® Ultra™ RNA Library Prep Kit for Illumina® (New England Biolabs, MA, USA) following manufacturer’s recommendations and index codes were added. The library quality was assessed on the Qubit@ 2.0 Fluorometer (Thermo Fisher Scientific, MA, USA) and Agilent Bioanalyzer 2100 system (Agilent Technologies, Waldbron, Germany). At last, the library was sequenced on an IlluminaHiseq2500 platform and 250 bp paired-end reads were generated (Guangdong Magigene Biotechnology Co., Ltd. Guangzhou, China).

### 2.5. Bioinformatic analysis of the sequencing data

Paired sequences were joined, and the adaptors, barcodes and primers were trimmed to remove low quality sequences and gained clean reads. Clean reads were mapped through the overlap relationship between PE reads into tags to obtain clean tags, and then the tags were clustered into OTU at 97% similarity. The OTU -based analysis of the alpha diversity indices, including the Shannon index, Chao 1, coverage, dominance and observed-species was performed with Mothur, and were calculated by using the QIIME platform (http://qiime.org/). A metagenomic biomarker discovery approach was employed with LEfSe (Linear discriminant analysis) coupled with effect size measurement which performs a nonparametric wilcoxon sum-rank test to assess the effect size of each phylum (13). The sequence data were processed on the quantitative insights into Microbial Ecology platform according to <u>Caporaso *et al*</u> (14) and the RDP classifier identified the microbial taxonomic status of the OTU representative sequence.

### 2.6. Statistical analysis

The diversity of samples was calculated to obtain the species richness and uniformity of samples, and the common and unique OTU of different samples or different groups could be known by Venn diagram and petal diagram information with R software. Three non-parametric tests were employed to determine the differences in the microbial structure between different cropping years, and heatmap and histogram based on relative abundance of phylum were produced with a heatmap package. Phylogenetic trees were constructed based on OTU representative sequences, and the community structure differences of different samples and groups could be further compared through sequencing analysis of PCoA and NMDS combining species abundance information. Correlation test and variation partitioning analysis (VPA) were performed by R software based on the key genus (relative abundance ≥ 1%). Pearson or spearman correlation index between species or samples based on the otu_table was calculated by R software and visualized by sytoscape (http://cytoscape.org/). Principal components analysis (PCA) was analysed based on the otu_table of the relationship between microbial biomass composition and the environmental factors was carried out by using the CANOCO5 software.

## 3. Results

### 3.1. Microbial community abundance and diversity

Fungal and bacterial diversity, richness and observed-species were higher in the rhizosphere soil compared with those in the unplanted soil. Shannon index in soils treated with chemical fertilizer only was more than that in soils treated with furfural residue for fungi in fields with a planting history longer than 20 years and for bacteria in all the fields. Chao 1 and observed-species in furfural residue-treated soils were more than those of chemical fertilizer-treated soils. In addition, microbes dominated more in unplanted soil than in the rhizosphere soil (Table 1).

**Table 1.**
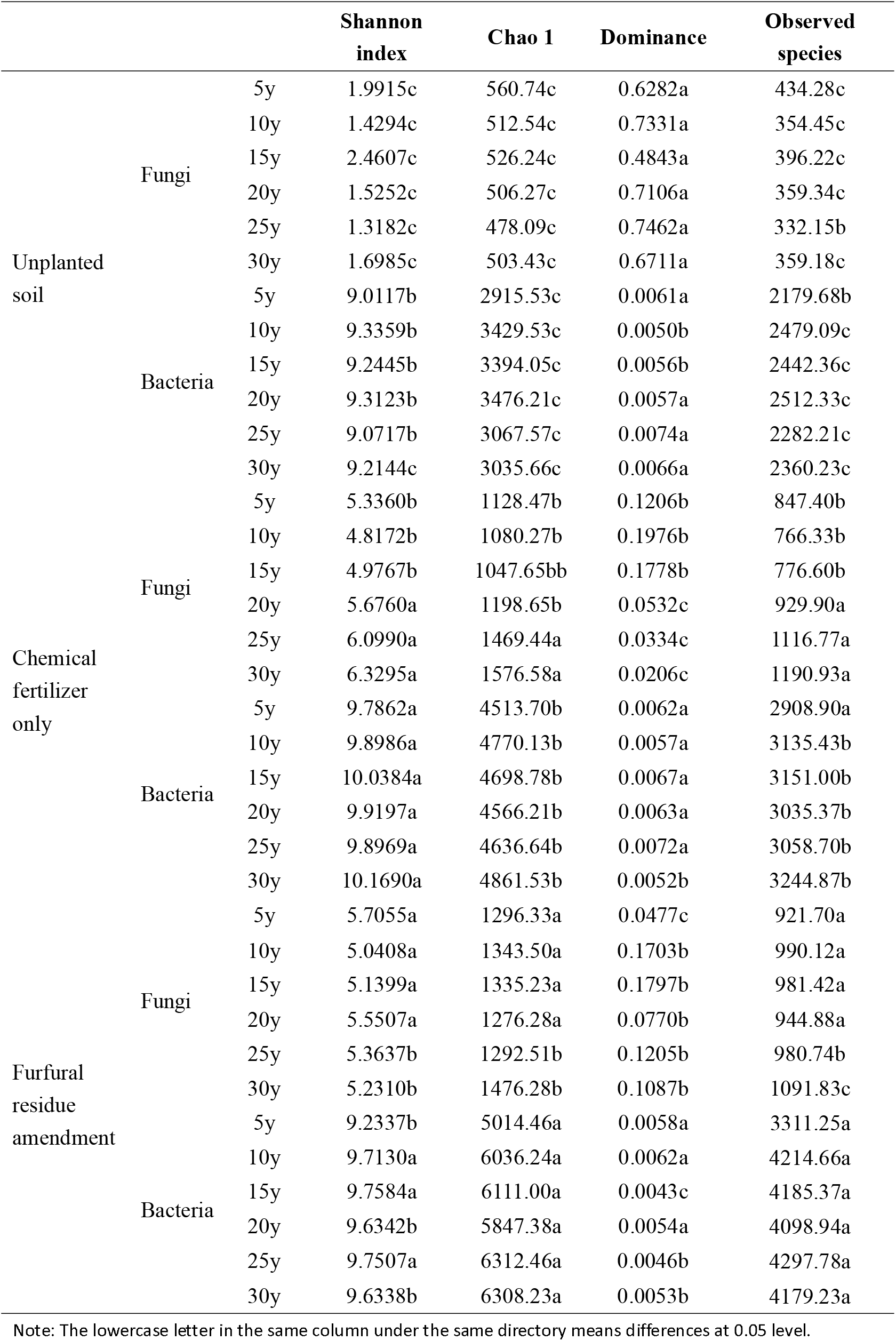
Shannon index, Chao 1, dominance, and observed species of fungi and bacteria of rhizosphere soil of different treatments

Venn diagram showed that 159 fungal OTUs were common among different years of continuous cropping in unplanted soil (Fig. 1A), while 172 were common among chemical fertilizer-treated soils (Fig. 1B). A total of 530 fungal OTUs were common among the furfural residue treatments. Unique fungal OTUs of the rhizosphere soil were more than those of the unplanted soil with increase in years of continuous cropping (Fig. 1). A similar difference was observed in both common and unique bacterial OTUs of the rhizosphere soil (Supplementary Fig. S1A–S1C). A significant increase in the microbial community diversity was observed, and more OTUs were induced in the rhizosphere soil. The species richness increased after planting and fertilization.

**Fig. 1A.**
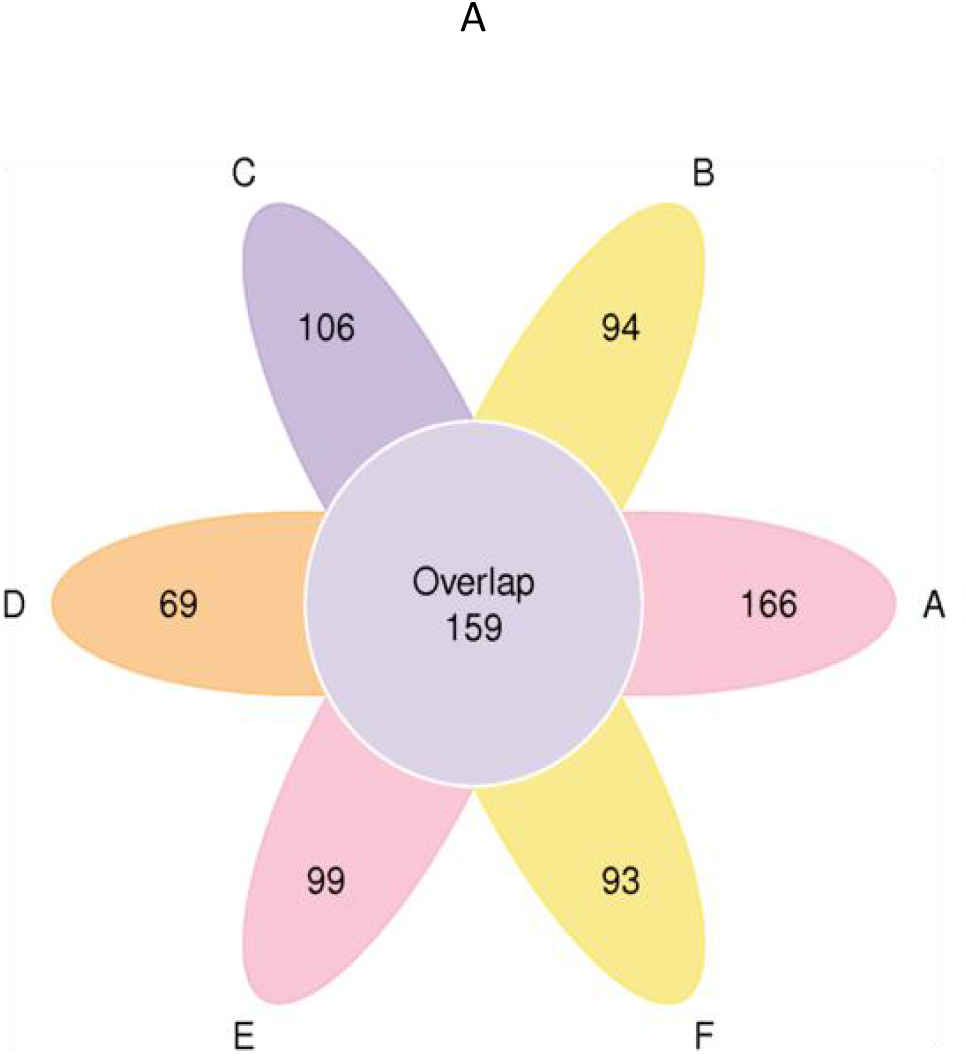
Petal graph of fungi OTUs presented in unplanted soil. The uppercase of A, B, C, D, E and F represents continuous cropping of corn seed production of 5, 10, 15, 20, 25, and 30 years, respectively.

**Fig. 1B.**
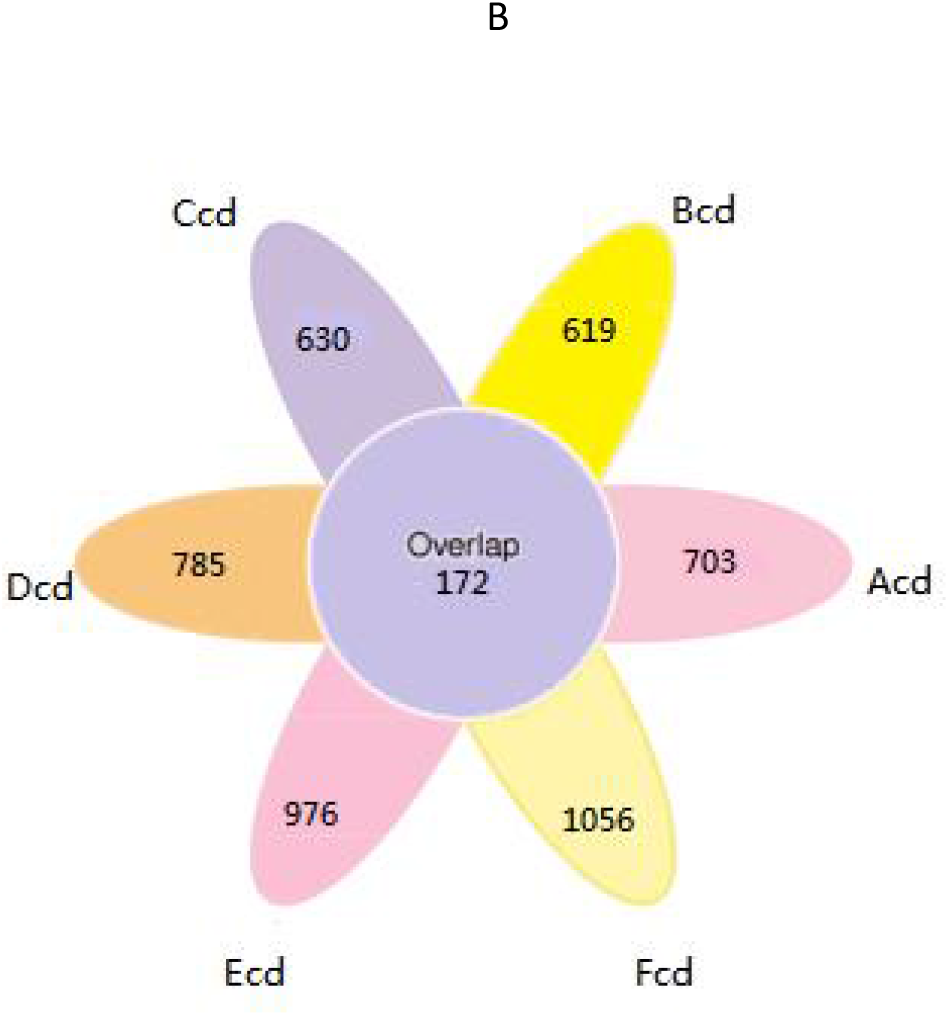
Petal graph of fungi OTUs presented in chemical fertilizer only amended soil. The lowercase of cd means chemical fertilizer only at bell mouth stage. The uppercase of A, B, C, D, E and F represents continuous cropping of corn seed production of 5, 10, 15, 20, 25 and 30 years, respectively.

**Fig. 1C.**
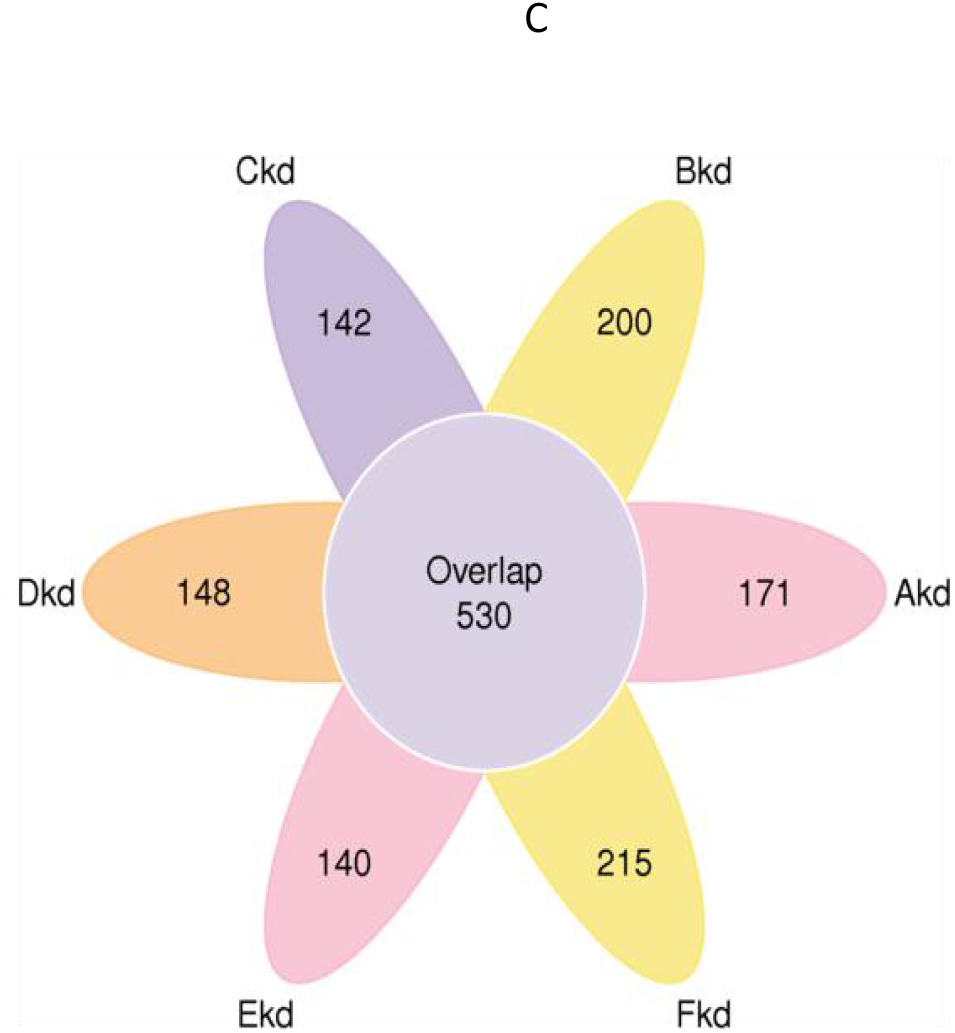
Petal graph of fungi OTUs presented in the furfural residue amended soil. Th lowercase of kd means furfural residue amendment at bell mouth stage. The uppercase of A, B, C, D, E and F represents continuous cropping of corn seed production of 5, 10, 15, 20, 25 and 30 years, respectively.

### 3.2. Composition of the fungal microbiota of rhizosphere soil

#### 3.2.1 Phylum composition of the fungal community in rhizosphere soil

A total of 101,307, 211,144, and 206,009 raw tags were analyzed and were grouped into 98,765, 204,295 and 198,278 ITS sequence from samples of seasonal soil batches. Approximately, 99.21% of the reads were utilized for metagenomic contig construction. Significant differences in fungal phyla were observed between the soil treated with and without amendment with furfural residue. Ascomycota and Mortierellomycota were dominant in soils treated with chemical fertilizer only, while Ascomycota and Basidiomycota were dominant in soil treated with furfural residue (Fig. 2A). The relative abundance of Basidiomycota increased while that of Ascomycota decreased in the soil treated with furfural residue with increase in years of continuous cropping (Fig. 2A).

**Fig. 2A.**
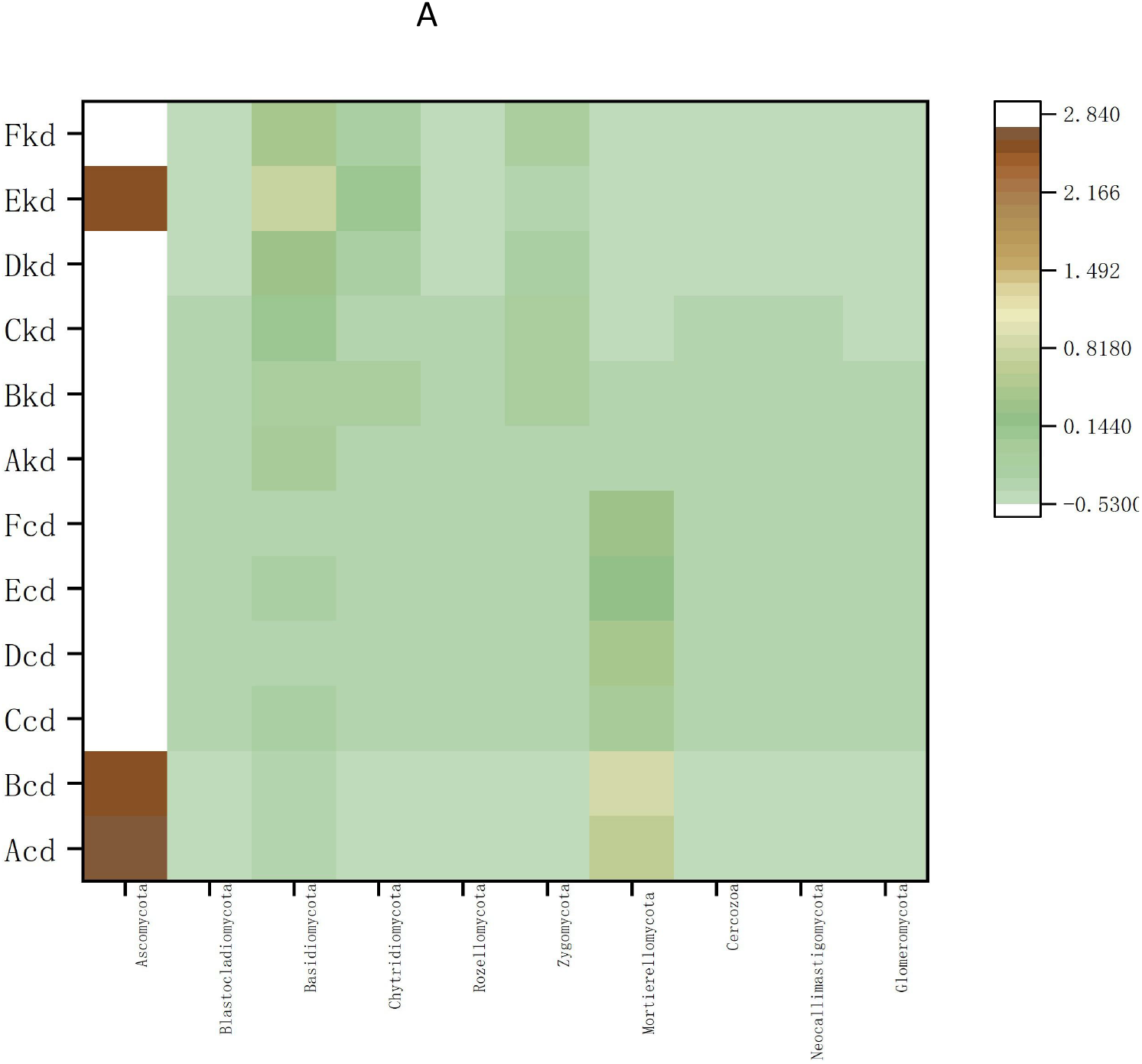
Heatmap of fungal phylum of rhizosphere soil at bell mouth stage. The lowercase of kd means furfural residue amendment at bell mouth stage, cd means chemical fertilizer only amendment at bell mouth stage.The uppercase of A, B, C, D, E and F represents continuous cropping of corn seed production of 5, 10, 15, 20, 25 and 30 years, respectively.

**Fig. 2B.**
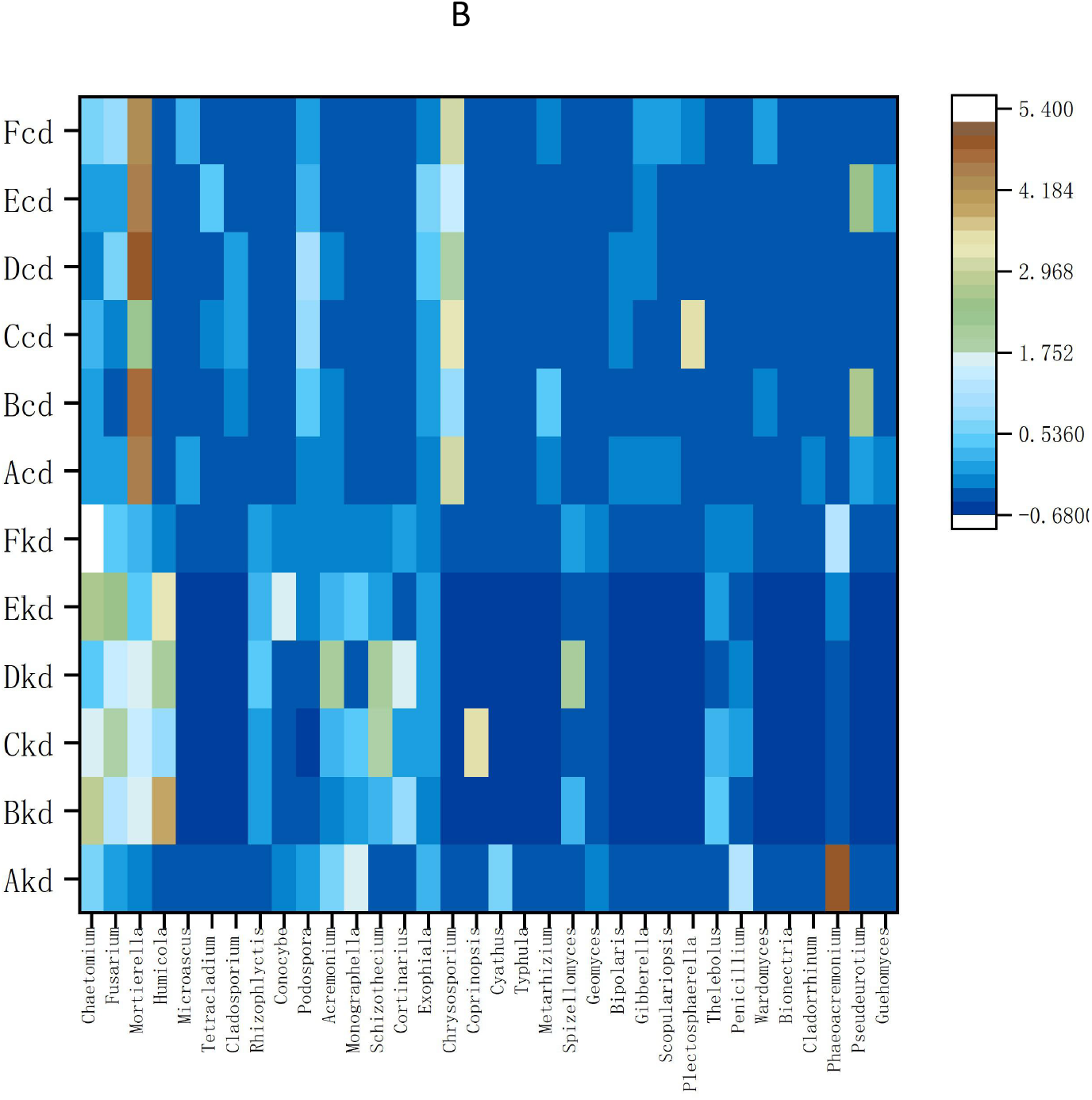
Heatmap of fungal genus (B) of rhizosphere soil at bell mouth stage. The lowercase kd means furfural residue application at bell mouth stage, cd means chemical fertilizer application at bell mouth stage. The uppercase of A, B, C, D, E, and F represents continuous cropping of corn seed production of 5, 10, 15, 20, 25 and 30 years, respectively.

Rhizosphere fungal microbiota from furfural residue-treated soil was significantly different from that retrieved from unplanted soil, and the richness of dominant phyla differed significantly among the seasonal soil batches. In unplanted soil, Ascomycota and Zygomycota were the two dominant phyla, while Chytridiomycota and Glomeromycota were least abundant (Supplementary Fig. S2A). In the rhizosphere of furfural residue-treated soils, Ascomycota, Basidiomycota and Chytridiomycota were dominant, while Zygomycota was least abundant (Supplementary Fig. S2B –S2C). A new inhabitant was the phylum Neocallimastigomycota, which increased with increase in continuous cropping years in the soils treated with furfural residue. Streptophycophyta was occasionally detected in the seasonal soil batches (Supplementary Fig. S2B), Fungi_ phy_ Incertae _ sedis was occasionally presented atthe bell mouth stage (Supplementary Fig. S2C).

#### 3.2.2 Genus composition of fungal community in rhizosphere soil

The community composition and diversity of fungal genera differed significantly between chemical fertilizer-treated soils and furfural residue-treated soils. *Chaetomium, Humicola, Monographella* and *Cortinarius* were more in the soils treated with furfural residue, while *Mortierella, Chrysosporium* and *Podospora* were more in the soils without furfural residue treatment (Fig. 2B). The top 30 fungal genera found in soils treated with or without furfural residue were totally different except for *Chaetomium, Fusarium, Podospora* and *Acrimonium*. The community composition of fungal genera shifted distinctly, and the microbial structure of rhizosphere soil was reestablished. The potentially pathogenic *Alternaria, Trichocladium, Bipolaris, Solicoccozyma, Gibberella* and *Cladosporium* were not detected, while the beneficial *Penicillium, Schizothecium* and *Rhizophlyctis* were present in the rhizosphere of soils treated with furfural residue (Fig. 2B).

Similarly, the community composition and diversity of fungal genera differed significantly among the treatments over the seasonal soil batches. *Alternaria, Trichocladium, Gibberella, Ciliophora, Candida, Solicoccozyma, Ilyonectria, Pseudogymnoascus, Metarhizium* and *Clonostachys* were specific inhabitants in the unplanted soil (Fig. 3A), while *Mycosphaerella, Camarops, Cercophora* and *Cryptococcus* were prevalent during seedling stage. *Fusarium* increased with increase in continuous cropping year at seedling stage (Fig. 3B). *Mortierella, Spizellomyces, Trichocladium* and *Verticillium* dominated the unplanted soil, *Typhula, Staphylotrichum, Coprinopsis* and *Ustilago* were new inhabitants in the rhizosphere soil at bell mouth stage. *Penicillium* in the rhizosphere soil increased within 5 to 20 years of continuous cropping; however, it gradually depleted from the rhizosphere soil after 20 to 30 years of continuous cropping (Fig. 3C). The genus *Spizellomyces* was detected in the soil in March and July; *Myrothecium* and *Ophiosphaerella* were dominant in the rhizosphere soil in March and May, while *Humicola, Rhizophlyctis, Cortinarius, Acremonium, Cyathus, Madurella, Phaeoacremonium, Geomyces*, and *Sarocladium* were dominant in the rhizosphere soil in May and July (Fig. 3). The increase during the seedling stage was notable.There was a significant decrease in the genera *Fusarium, Humicola, Chaetomium, Microdochium* and *Monographella* during the bell mouth stage. Pathogenic *Alternaria, Trichocladium, Gibberella* and *Solicoccozyma* disappeared in May and July after the application furfural residue. The relative abundances of *Aspergillus and Verticillium* decreased with plant growth. The relative abundance of *Mortierella, Conocybe* and *Schizothecium* in rhizosphere soil decreased at the seedling stage, and then increased at the bell mouth stage (Fig. 3).

**Fig. 3A.**
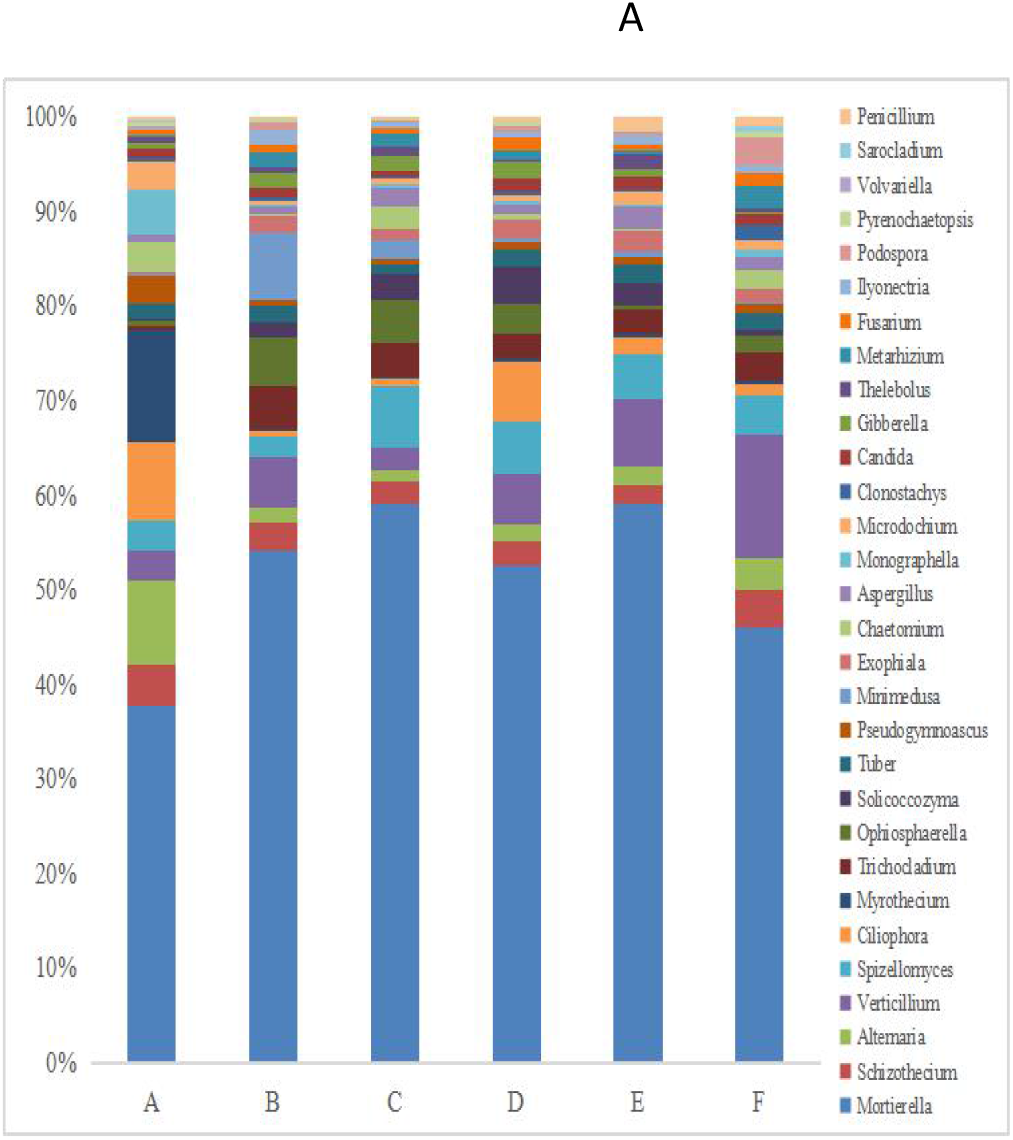
Genus of rhizosphere fungi of soil in March (unplanted soil). The uppercase of A, B, C, D, E and F represents continuous cropping of corn seed production of 5, 10, 15, 20, 25 and 30 years, respectively.

**Fig. 3B.**
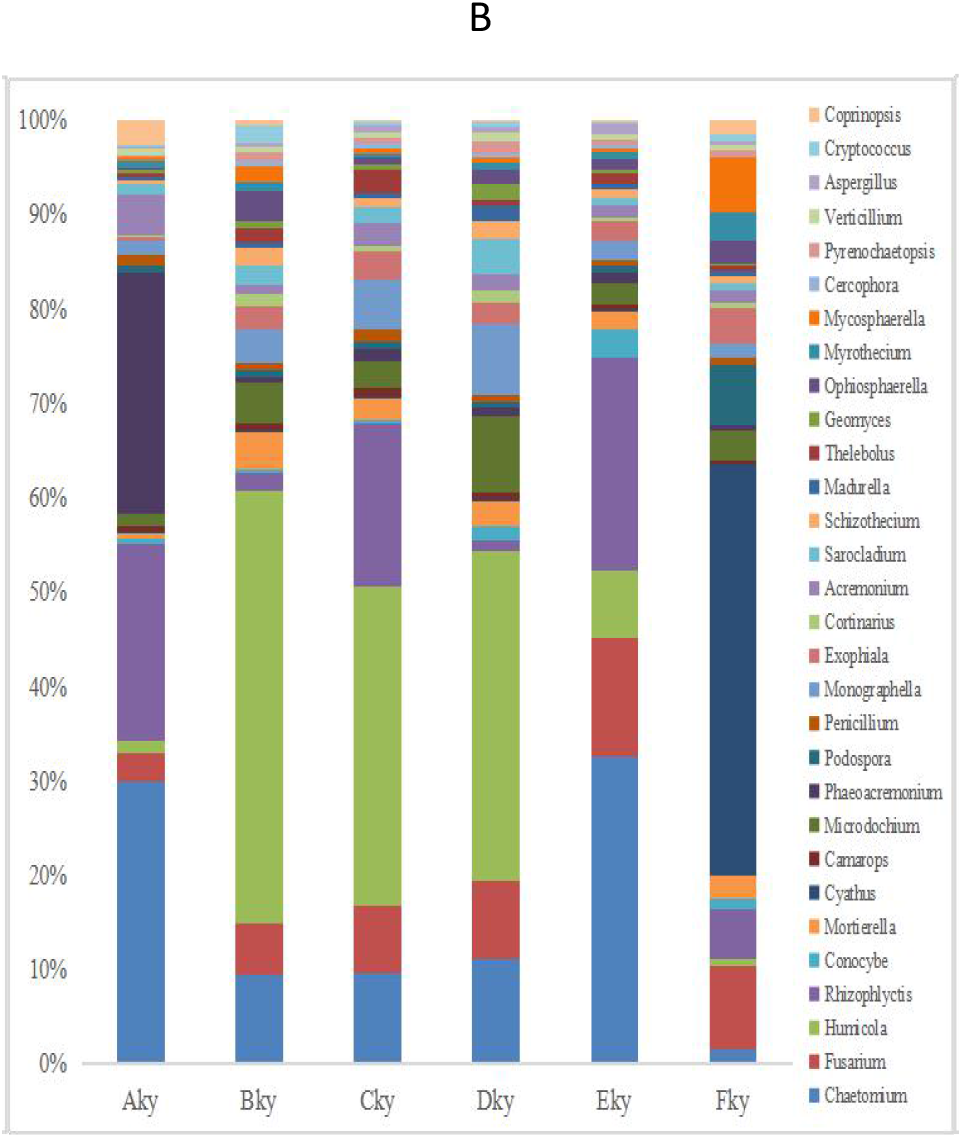
Genus of rhizosphere fungi of soil in May (the soil at seedling stage). The lowercase of ky means furfural residue application at seedling stage. The uppercase of A, B, C, D, E and F represents continuous cropping of corn seed production of 5, 10, 15, 20, 25 and 30

**Fig. 3C.**
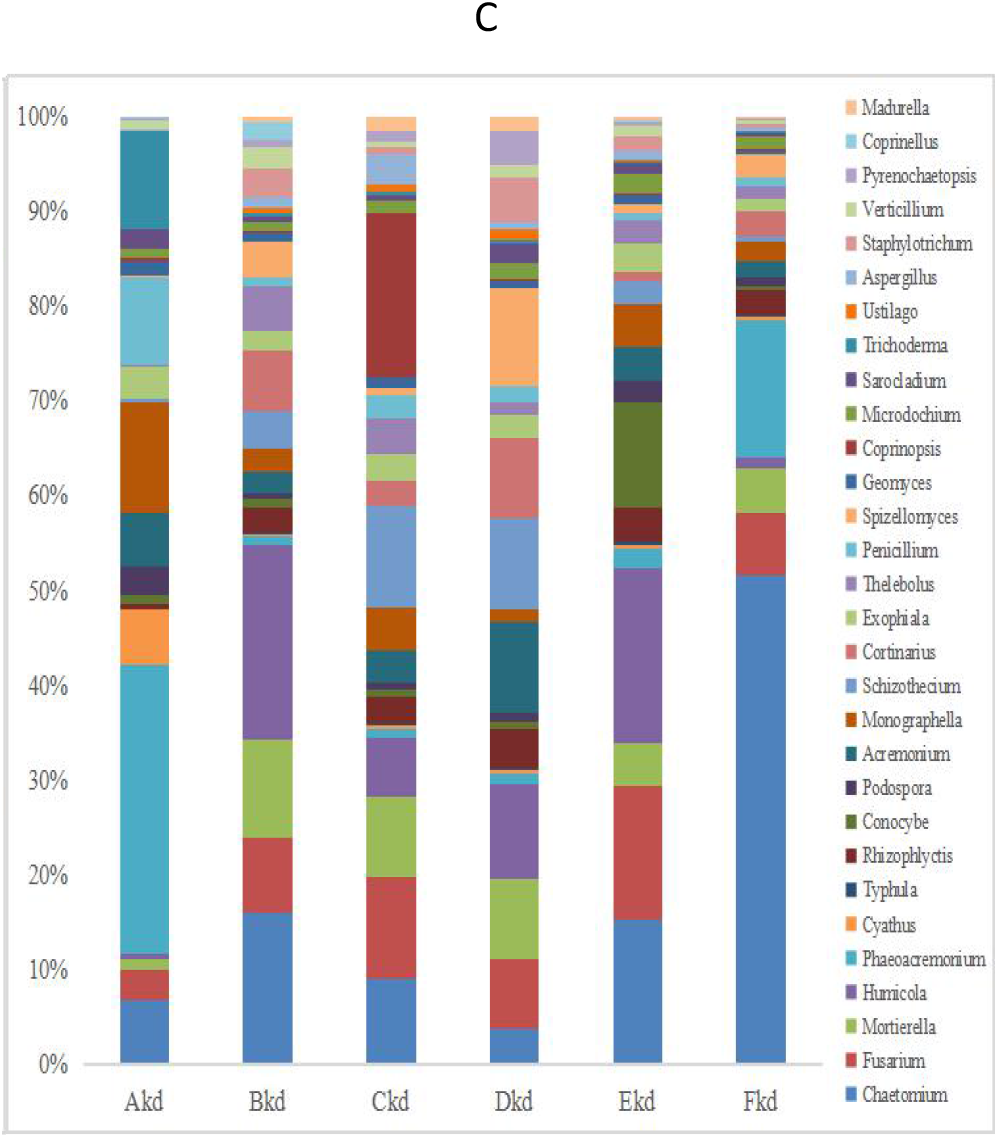
Genus of rhizosphere fungi of soil in July (the soil at bell mouth stage). The lowercase of kd means furfural residue application at bell mouth stage. The uppercase of A, B, C, D, E and F represents continuous cropping of corn seed production of 5, 10, 15, 20, 25 and 30 years,

### 3.3. Composition of bacteria communication in rhizosphere soil

#### 3.3.1 Phylum composition of the bacteria community in rhizosphere soil

A total of 91,973, 92,190, and 242,628 raw tags were analyzed and the sequences were grouped into 68,626, 69,756, and 162,042 16S sequences. The bacterial community in the rhizosphere of unplanted soil had smaller species richness and was greater evenness compared with that in the rhizosphere furfural residue treated-soils (Table 1).

The 16S rRNA gene sequences retrieved from the rhizosphere soil under continuous cropping were affiliated with 30 bacteria phyla and corresponded to 1,269 species-level OTUs. Actinobacteria and Thaumarchaeota in the rhizosphere decreased with furfural residue treatment and increased with chemical fertilizer treatment (Fig. 4A). In the rhizosphere soil, Acidobacteria and Bacteroidetes were dominant with furfural residue application, while Firmicutes and Chloroflexi were dominant with chemical fertilizer treatment. Rokubacteria and Gemmatimonadetes were less abundant in the soils treated with chemical fertilizer only compared with soils treated with residue. Proteobacteria, Bacteroidetes and Acidobacteria were the three dominant bacterial phyla under furfural residue treatment over different continuous cropping years (Fig. 4A).

**Fig. 4A.**
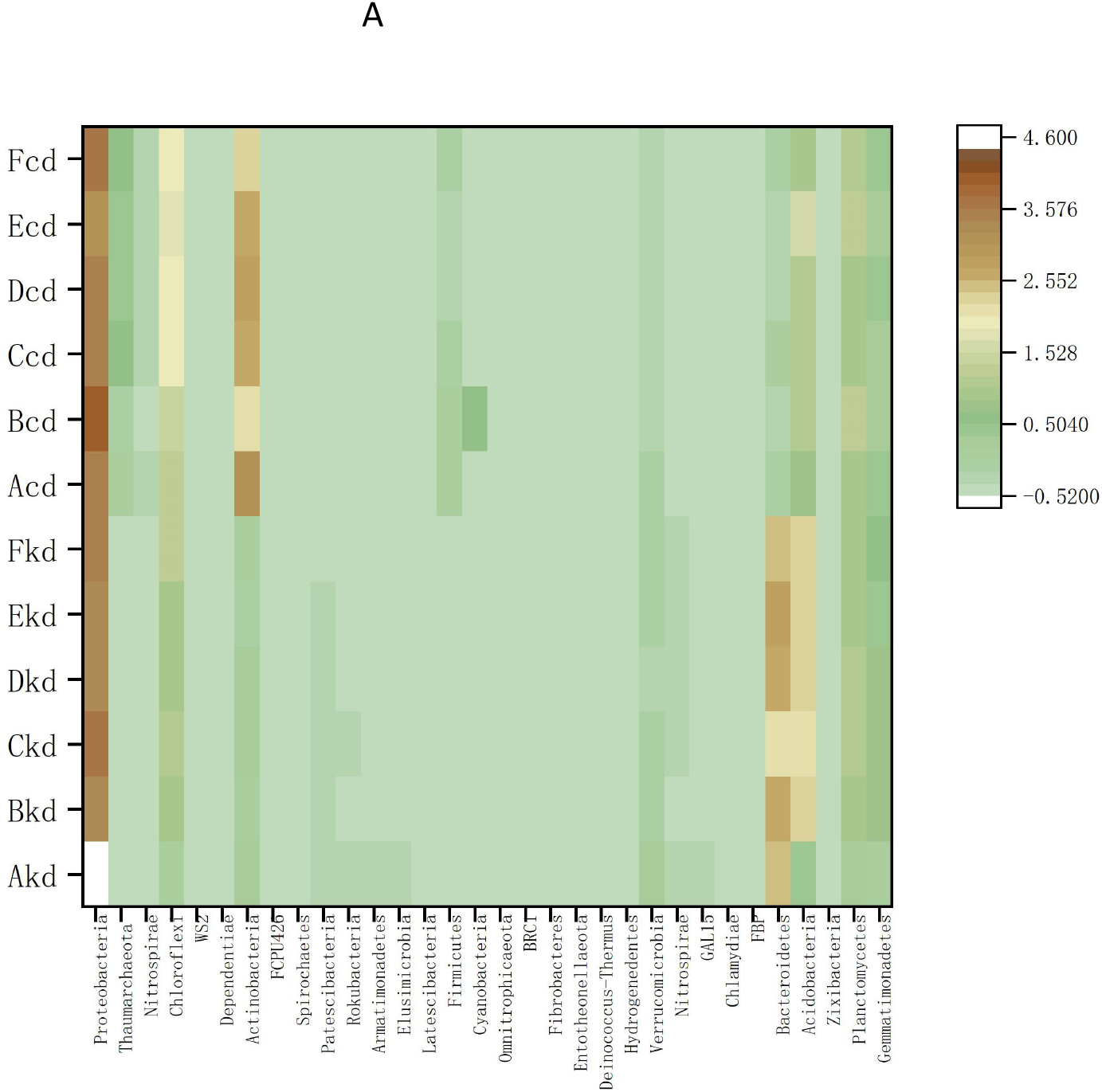
Heatmap of phylum of rhizosphere bacteria at bell mouth stage.The lowercase of kd means furfural residue application at bell mouth stage, cd means chemical fertilizer only application at bell mouth stage. The uppercase of A, B, C, D, E and F represents continuous cropping of corn seed production of 5, 10, 15, 20, 25 and 30 years, respectively.

**Fig. 4B.**
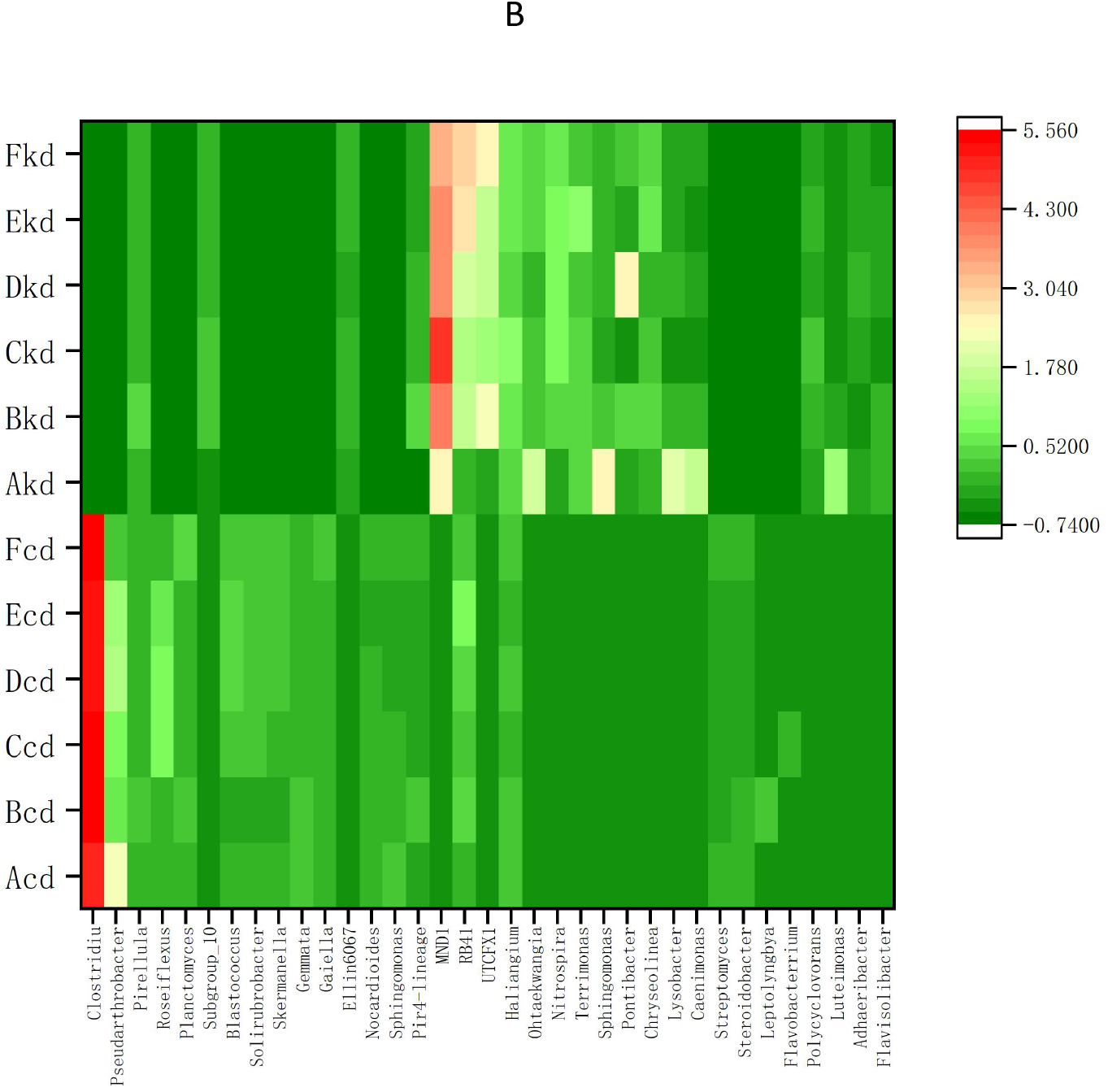
Heatmap of genus of rhizosphere bacteria at bell mouth stage. The lowercase of kd means furfural residue application at bell mouth stage, cd means chemical fertilizer only application at bell mouth stage. The uppercase of A, B, C, D, E and F represents continuous cropping of corn seed production of 5, 10, 15, 20, 25 and 30 years, respectively.

In the seasonal samples, phylum WPS-2 was prevalent in March and May; FCPU426 was specific to May and July; and Epsilonbacteraeota and Spirochaetes were specific to March and July. Proteobacteria and Bacteroidetes were abundant in May and sparse in July, while Acidobacteria showed an opposite trend. The bacterial genera Cyanobacteria, Firmicutes, Actinobacteria and Euryarchaeota depleted in unplanted soil and seasonal soil batches (Supplementary Fig. S3A–S3C).

#### 3.3.2 Genus composition of the bacterial community of rhizosphere soil

The relative abundance of *MND1, RB41, UTCFX1, Nitrospira, Ellin6067, Adhaeribacter, Flavisolibacter, Polycyclovorans* and *Terrimonas* was greater in furfural residue treated soils, while the abundances of *Clostridium, Pseudarthrobacter* and *Roseiflexus* was lesser(Fig. 4B).

In the seasonal soils, *UTCFX1, RB41*and *Terrimonas* were the dominant genera in March (Fig. 5A), while *MND1, RB41* and *UTCFX1* were dominant in May and July (Fig. 5B – C). *Arcticibacter, ADurb*.*Bin063-1, Stenotrophomonas, UTBCD1* and *Dongia* were present only in

**Fig. 5A.**
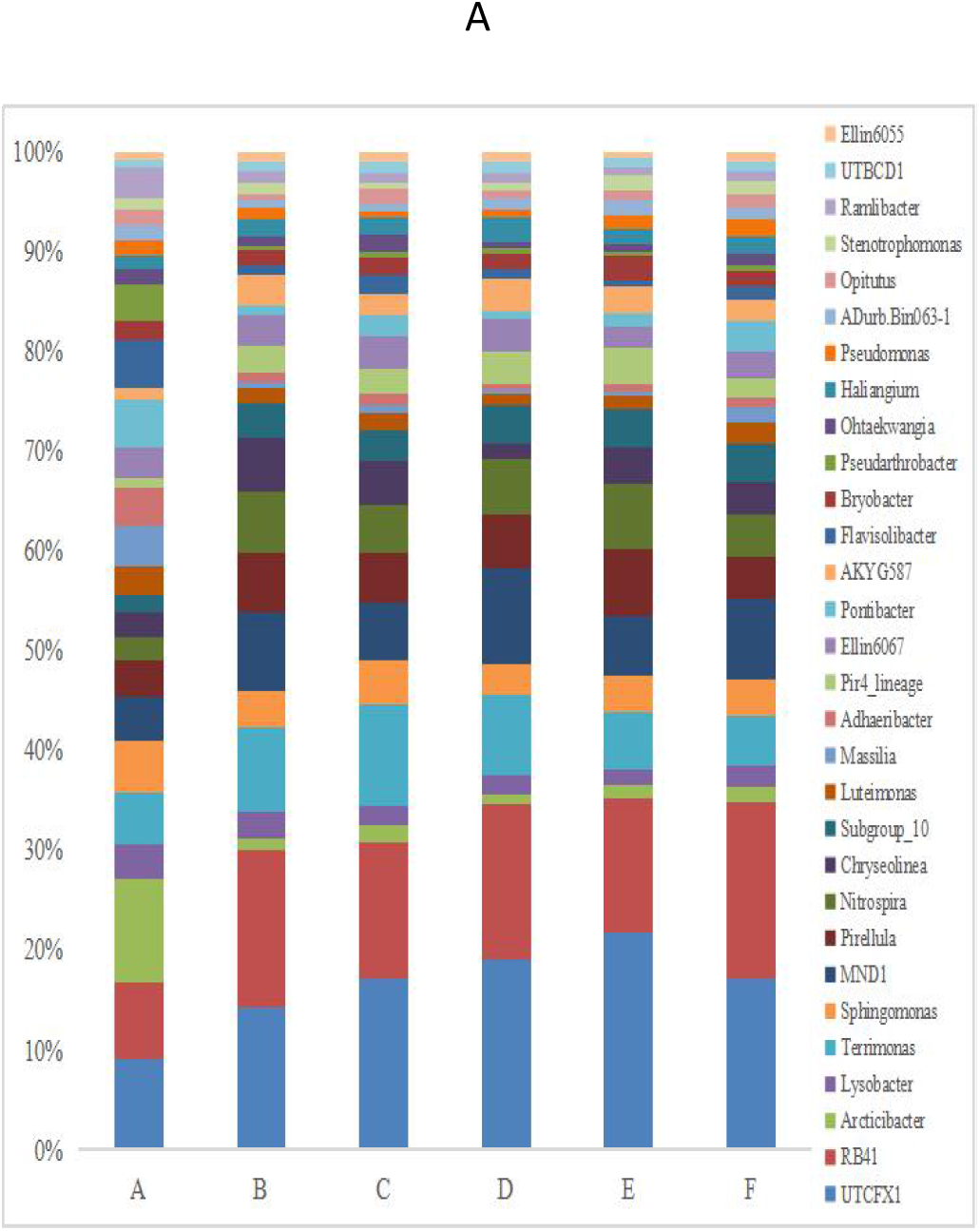
Genus of rhizosphere bacteria of soil in March (unplanted soil). Letter A, B, C, D, E and F represents continuous cropping of corn seed production of 5, 10, 15, 20, 25 and 30 years, respectively.

**Fig. 5B.**
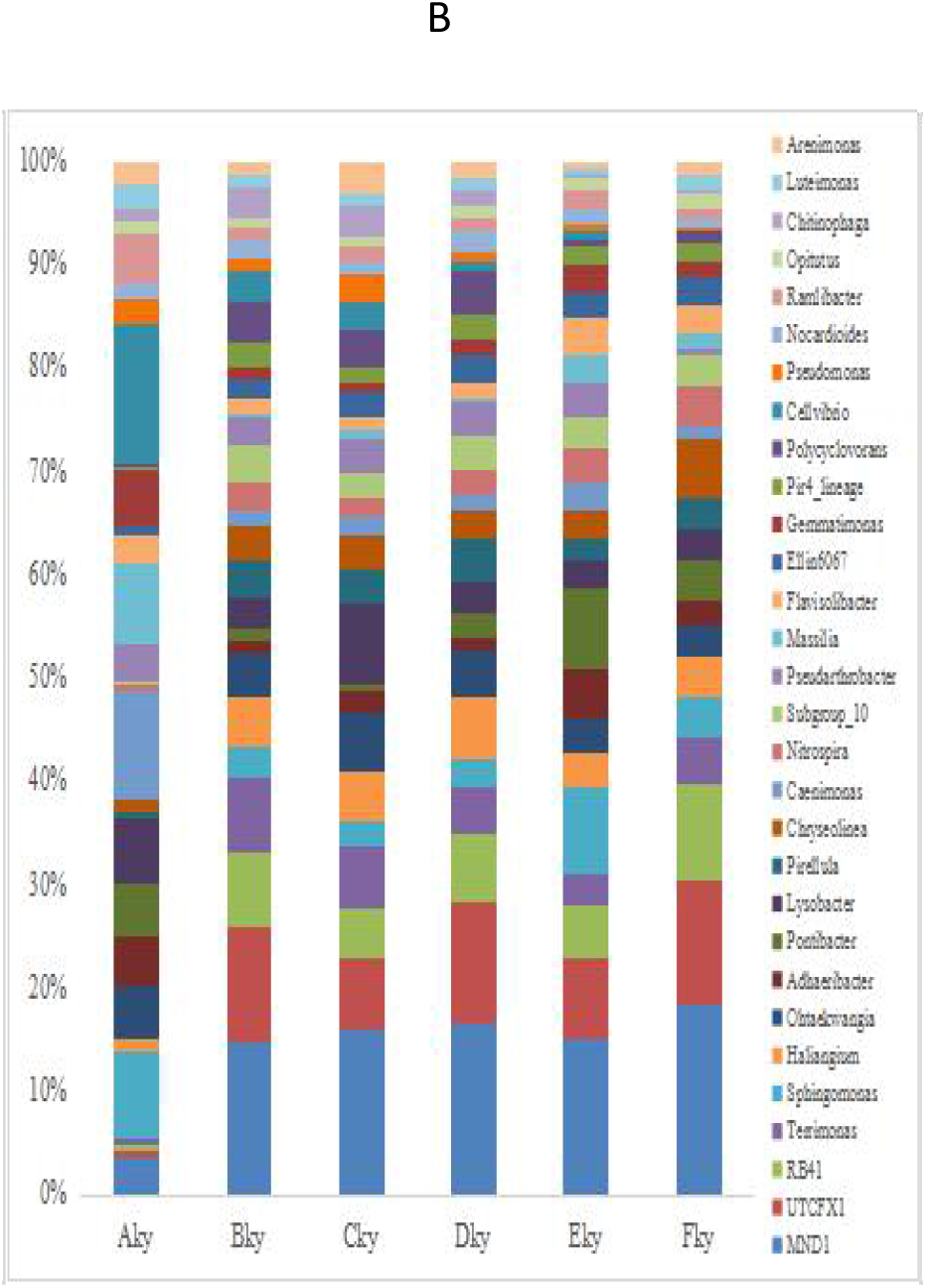
B Genus of rhizosphere bacteria of soil in May (seedling stage). The letter ky means furfural residue application at seedling stage. Letter A, B, C, D, E, and F represents continuous cropping of corn seed production of 5, 10, 15, 20, 25 and 30 years, respectively.

**Fig. 5C.**
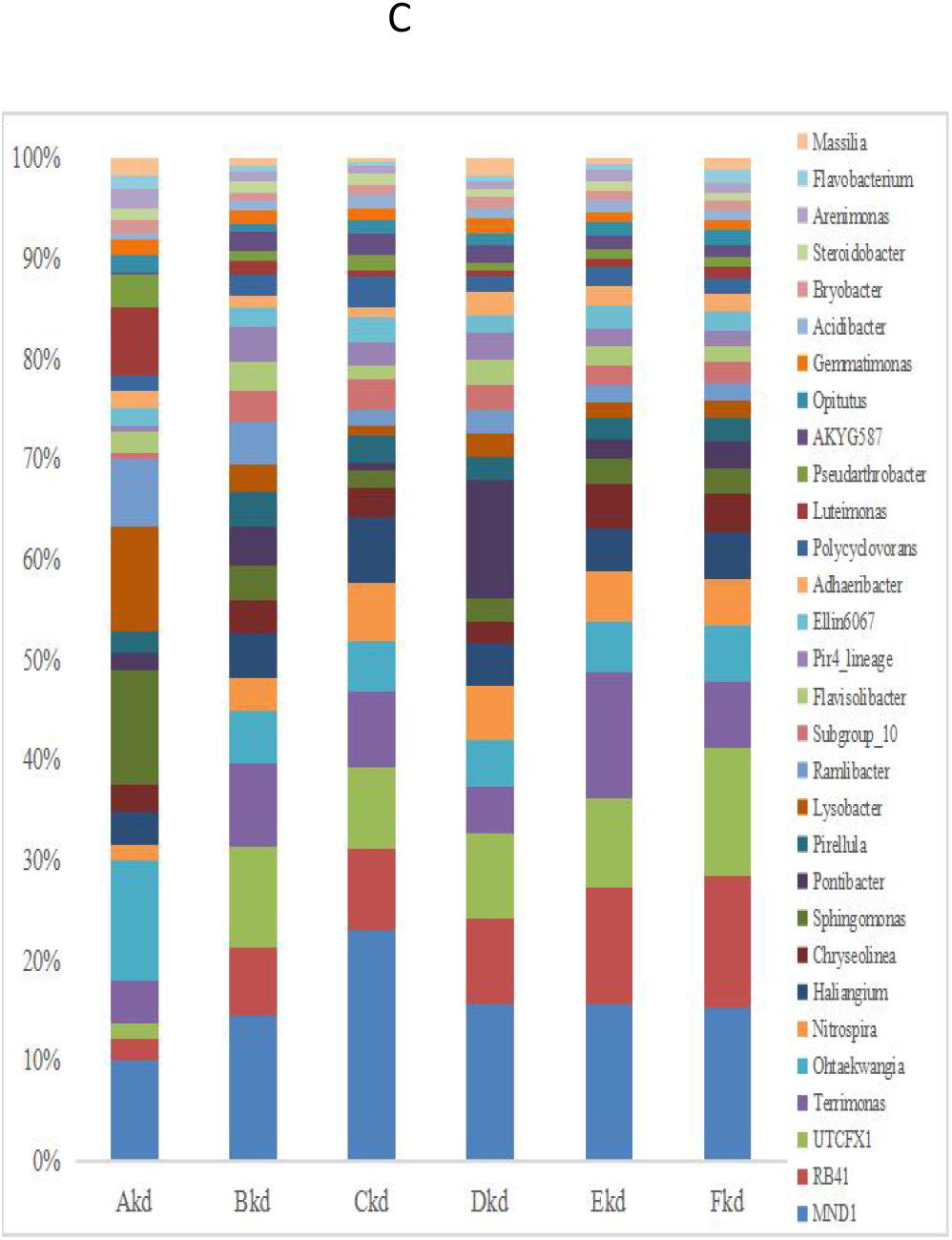
Genus of rhizosphere bacteria of soil in July (bell mouth stage). The letter kd means furfural residue application at bell mouth stage. Letter A, B, C, D, E, and F represents continuous cropping of corn seed production of 5, 10, 15, 20, 2B5 and 30 years, respectively.

March (Fig. 5A), while *Caenimonas, Cellvibrio, Polycyclovorans, Gemmatimonas*, and *Arenimonas* genera were restricted to the rhizosphere soil in May and July (Fig. 5B– C). *RB41, Terrimonas, Nitrospira, Pir4_lineage, AKYG587* first decreased and then increased with time, while *Lysobacter, Adhaeribacter, Massilia* and *Pseudarthrobacter* showed an opposite trend. The relative abundance of *Haliangium* and *MND1* increased, while those of *Pirellula, Ellin6067* and *Luteimonas* decreased in the seasonal soil batches (Fig. 5).

### 3.4. PCoA analysis

To examine the potential role of continuous cropping, fertilization and growth stage on the root-inhabiting fungal and bacterial assemblages, we compared the fungal and bacterial profiles obtained from the unplanted and planted soil samples with or without furfural residue application and at different growth stages (Fig. 6). Taken together, they explained 97.45% of the total variability in fungal community at the phylum level in the rhizosphere soil and 37.25% in bacterial community in the rhizosphere soil. Apparent clustering was due to the variability with fertilization and growth stage. Fertilizer treatment and seasonal soil batches together explained 96.36% variability in fungal communities. The first two axes of the PCA plot explained 94.09% of the total variability in data (Fig. 6A). Community diversity metrics clearly demonstrate that communities from the same treatments were similar and those from other treatments were distinct. The soil microbial community diversity after different continuous cropping years under the same treatment differed from each other; however, occasional overlap was observed between the samples. The variation of microbes in soils treated with furfural residue at different growth stages were similar and situated in the same quadrant, which indicates minimal effect of seasonal soil batches on the microbial community structure. The dissimilarity in community diversity profiles among unplanted soils, furfural residue treatments and chemical fertilizer only treatments is notable and apparent from the PCA ordination. Communities from unplanted soils and chemical fertilizer only treatments showed the maximum difference. The contribution of selected properties followed the trend of fertilizer > furfural residue > planting > seasonal soil batches. These results indicate that the microbial community structure is closely associated with fertilization (Fig. 6A). These variables together explained the variance in bacterial communities (Fig. 6B). The first two axes of the RDA plot explained 20.51% and 14.55% (35.06% in total) of the total variability. A significant correlation was observed between the microbial community diversity and the variations in bacterial community. Fertilization was the main factor that shaped the microbial community structure (Fig. 6B).

**Fig. 6A.**
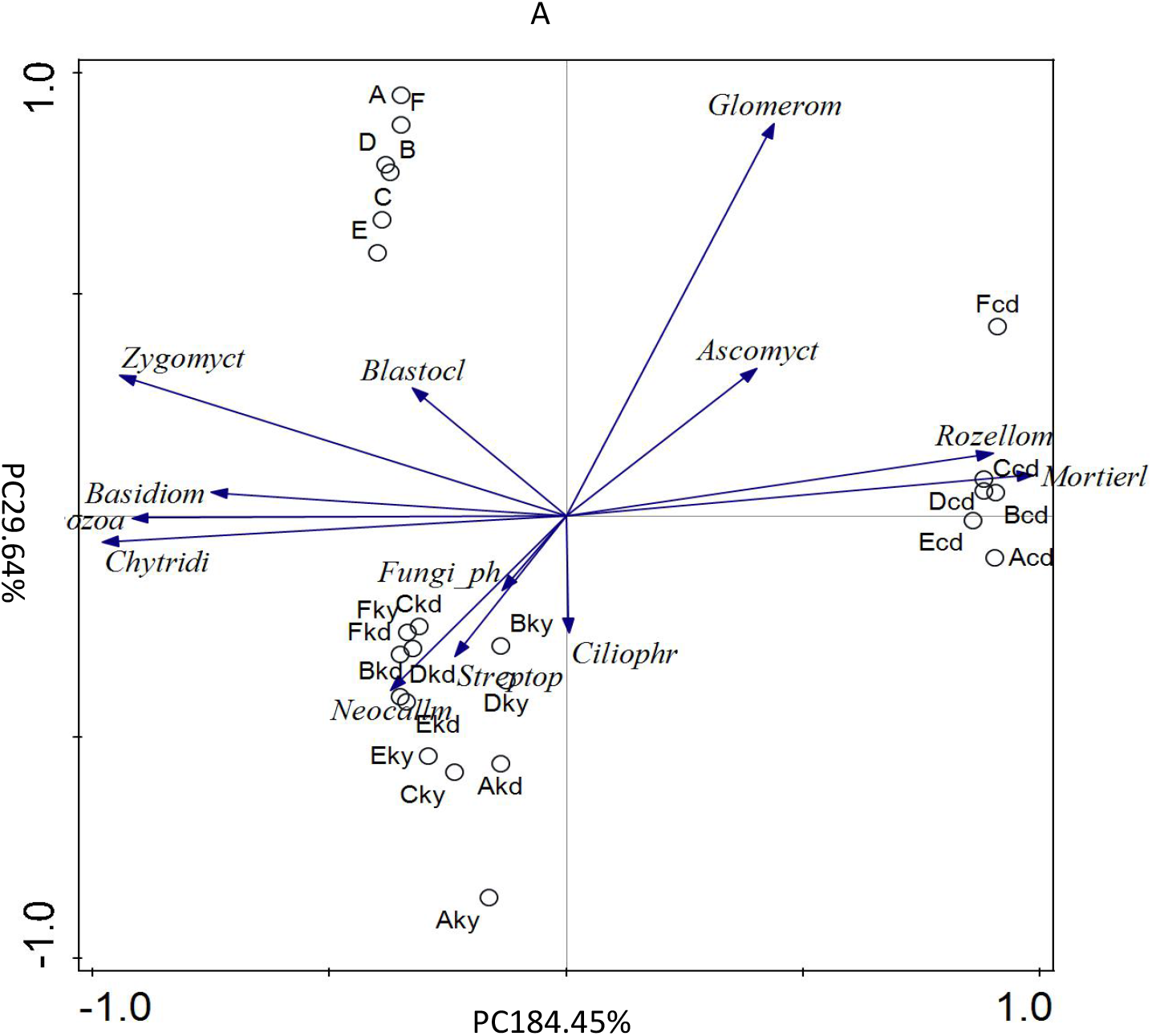
PCA analysis of the relationship between fungi phylum and different continuous cropping years. The lowercase of ky means furfural residue application at seedling stage, kd means furfural residue application at the bell mouth stage, and cd means chemical fertilizer only at the bell mouth stage. The uppercase of A, B, C, D, E and F represents continuous cropping of corn seed production in 5, 10, 15, 20, 25 and 30 years, respectively.

**Fig. 6B.**
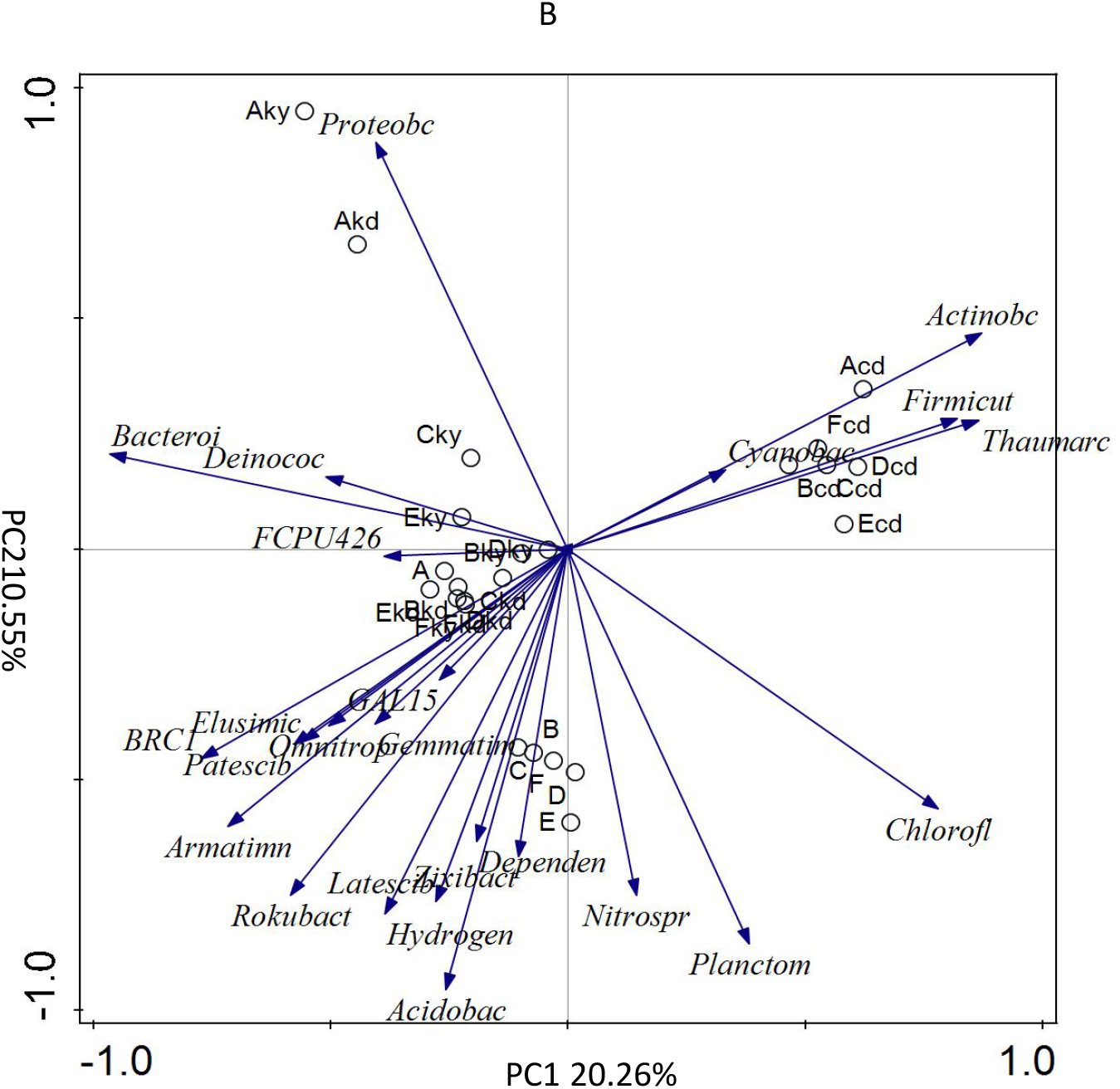
PCA analysis of the relationship between bacteria phylum and different continuous cropping years. The lowercase of ky means furfural residue application at the seedling stage, kd means furfural residue application at the bell mouth stage, and cd means chemical fertilizer only at the bell mouth stage. The uppercase of A, B, C, D, E and F represents continuous cropping of corn seed production in 5, 10, 15, 20, 25 and 30 years, respectively.

## 4. Discussion

The differences in the diversity and relative abundance of rhizosphere microbiota were investigated under different years (5–30) of continuous cropping with different fertilizers and at different growth stages. Rhizosphere microbiota was different among all soils (unplanted, chemical fertilizer only and furfural residue treated). The community divergence of unplanted soil under different years of continuous cropping was large and similar to that of soils with chemical fertilizer only and furfural residue treatments. While the same species variation after different continuous cropping years was inconsistent and non-linearly dependent, the rhizosphere microorganisms demonstrated specific variation after different continuous cropping years. Therefore, variations of key genera under fixed environmental conditions over different years of continuous monocropping need to be identified. A coordinated regulation among the host plant, root microbiota and environment is essential for plant growth (15), which will in turn lead to microbial division and diversity over different years of continuous cropping.

Maize is a staple crop infected by fungal, bacterial, and viral pathogens (16). Plant diseases caused by various pathogens are difficult to control. Understanding the variations in pathogenic organisms in the rhizosphere soil will help predict the occurrence of diseases and enable the use of efficient agents to control these epidemic diseases. Hafiz (17) reported seedling blight, stalk rot and leaf blight as the most important diseases of corn. The most prevalent diseases in the local area include rot disease, spot disease and wilt of corn. Studies have revealed rot disease and spot disease as most devastating in hot, humid tropical and temperate regions of the world. The dominant increase in *Fusarium, Alternaria, Trichocladium, Gibberella, Acremonium* and *Monographella* increase the probability of disease occurrence. *Fusarium* spp. in soil reduces the production and quality of plants (18). *Alternaria* species induce several diseases, such as early blight, black spot, brown patch, stem wilt and brown spot disease (19,20). Furfural residue application decreased the diversity of fungal microbiota, including major fungal pathogens. Dominant reduction in *Fusarium, Humicola, Chaetomium, Microdochium* and *Monographella* at bell mouth stage; disappearance of *Alternaria, Gibberella, Trichocladium* and *Solicoccozyma* and continuous decline in *Aspergillus* and *Verticillium* were observed during the entire life cycles. Correspondingly, corn root rot-related fungi, such as *Fusarium* and *Alternaria*, were either absent or less abundant over different years of continuous cropping following the application of furfural residue. Members affiliated with these decreasing fungal genera included plant pathogens, which may lead to plant disease occurrence under suitable conditions. The microbial populations and communities reestablished by furfural residue application improved soil health with increased resistance to disease occurrence. Therefore, furfural residue can be potentially used as a soil amendment in disease management. However, the appearance of potential pathogens such as *Acremonium* and *Ustilago* after amendment with furfural residue indicated only partial reduction in the number of soil pathogens.

Some of the non-pathogenic fungal and bacteria genera may act as potential biocontrol agents against fungal and nematode pathogens (21,22). Identification of beneficial rhizosphere microbes from maize fields treated with furfural residue will initiate studies. Studies have revealed that *Exophiala* and some *Collectotrichum spp*. have been suggested to promote plant growth via hormone production and absorption of phosphorus under abiotic stresses (23,24). *Pseudomonas* species suppress pathogens in the rhizosphere (25,26). *Trichoderma* spp. and *Bacillus* spp. were used to be control banana Fusarium wilt (27,28,29), and a combination of biocontrol agents with organic matter improved their biocontrol ability (30,31). The genus *Chaetomium* consists of 300 cellulolytic fungal species (32), and *Penicillium* are active producers of xylanolytic enzymes that degrade xylan, a major component of lignocellulose (33). *Neocallimastigomycota* plays an important role in the degradation of plant material (34). *Mortierella, Chaetomium, Conocybe, Cercophora, Coprinopsis* and *Schizothecium* were abundant at bell mouth stage in soils with fertilization. Many are affiliated with cellulose and lignin decomposition and degradation (35,32, 36). Acidobacteria has an important role in the carbon cycle (37) and Acidobacteriaceae family of this phylum has the ability to degrade simple carbon compounds and microbial polysaccharides, including cellulose (38). Bacteroidetes was found to play a pivotal role in the remineralization of high molecular weight organic matter (39). This phylum has been Gram-negative pioneers in decomposing organic matter (40). Firmicutes was the dominant bacterial genus that significantly increased during reductive conditions (32), and Chloroflexi isolates have been shown to scavenge atmospheric H_2_ produced by Actinobacteria and Acidobacteria (41). Webber et al. showed that Proteobacteria and Actinobacteria are capable of oxidizing atmospheric CO_2_ (42,43). This study showed that the microbes involved in carbon metabolism shifted from heterotrophic Proteobacteria and Actinobacteria to autotrophic Bacteroidetes. All these results obviously indicated that the reductive-like microbes in soil treated with chemical fertilizer only shifted to microbes involved in degradation and mineralization. Meanwhile, the microbial population and community were reestablished after furfural residue application. Some of the identified rhizosphere microbes can directly obtain amino acids from root exudates; amino acids are important nitrogen and/or carbon sources in this habitat (44). A few fungal and bacterial genera may be potential agents that could regulate nutrient absorption and regulate metabolism under adverse environment. The application of furfural residue increased the overall abundance of core microbes, including *MND1, Haliangium, RB41, UTCFX1, Nitrospira, Terrimonas, Cellvibrio, Flavisolibacter, Steroidobacter* and *Adhaeribacter* majority are involved in the metabolism of carbon, nitrogen, phosphorus and sulfur metabolism (45) and material degradation (46,47). These activities will contribute to soil nutrient cycling, plant growth improvement, material degradation and nutrient utilization. Nutrition absorption is another important factor related to the changes in rhizosphere microbiome (48). The application of furfural residue had a significant impact on the microbial subdivision of community composition. The microbial community composition in the rhizosphere changes with time and in response to the root exudation patterns that vary during plant growth and seasonal responses (49). In this study, furfural residue application promotes a beneficial soil environment.

Previous studies have shown that many abiotic factors, such as soil type and texture (50), soil pH (51), carbon availability (52) and biotic factors influence the microbial communities. Soil organic carbon content and soil type were the important abiotic factors that shaped the microbial communities (53). Furfural is a carbon source produced from corncobs, and it poses a serious threat to the local environment (54). This residue is rich in lignin and cellulose, and 2D-NMR (Two dimension NMR) and Py-GC/MS (Pyrolysis - gas chromatogram/mass spectrometer) analyses of the residue revealed the presence of lignin and depolymerized cellulose moieties; high content of phenol was produced in the process (55). The application of these residues will increase the amount of degradation-resistant carbon and aromatic materials in the soil, which will inevitably affect the microbes involved in resolving, releasing and reutilizing these materials. Previous studies also demonstrated that microorganisms were highly inactive in bulk soil and showed increased activities in the rhizosphere soil (56) particularly during the plant developmental stages (57). The differences in micro-environments related to root exudation may primarily have caused the variation of microbes. Present study shows that the microbial taxa recruited to the rhizosphere from the soil microbial reservoir vary among growth stages, and a given growth stage apparently selects a particular microbiome. The diversity of microbes in the bulk soil, which is the resource for the rhizosphere, is important since the effect of plant roots is often temporary (56). This identifies plant type as an important environmental variable that influences the quantitative and qualitative composition of root-inhabiting microbiota. Seasonal variations in the activity and relative abundance of rhizosphere microbial communities are dependent on plants (58). Same plant species with different microbiota in the fields of the current study indicates that unrelated microbial communities shift over time. Community components of the corn seed rhizosphere under planting conditions were similar despite the differences in continuous cropping years. This suggests a host-driven selection for particular traits. Only few bacterial taxa, such as Proteobacteria, Actinobacteria, Acidobacteria, Bacteroidetes and Chloroflexi, were abundant in both unplanted and rhizosphere soils. Shifts in the rhizosphere microbial communities may contribute to the differences in resistance to soil-borne disease under different plant root exudates. The composition of root microbiota may be influenced by soil type, which could reflect differences in the natural inocula. However, all fields of this study had sandy loam soils. The results indicate that a significant host-independent community change occurred in seasonal soil batches as well as host-dependent community shifts in rhizosphere soils treated with furfural residue. The shift and depletion in potential pathogens may lead to differences in disease occurrence and severity, which affects the type of rhizosphere microbes and the development of soil-borne plant diseases (59,60). These findings together indicate that the microbial community composition in rhizosphere changes over time and in response to plant growth and seasons. Future studies should aim to identify the microbes and test their functional roles as targets. Focus should be to identify potentially useful microbes from the population, with common or uncommon functions, selected from a given rhizosphere.

## 5. Conclusion

The community divergence was obvious after different years of continuous cropping, and the variation in a species after different years of continuous cropping was inconsistent and non-linearly dependent. Unplanted soil, chemical fertilizer-treated soil and furfural residue-treated soil recruited distinct rhizosphere microbiota under field conditions.The rhizosphere microorganisms demonstrated specific variation after different years of continuous cropping. Microbial community diversity increased, and many OTUs were induced in the rhizosphere soil than those in the unplanted soil. Ascomycota and Mortierellomycota were dominant in chemical fertilizer only treatment. Ascomycota and Basidiomycota were prevalent in furfural residue-treated soils, while Ascomycota and Zygomycota were dominant in the unplanted soil. Acidobacteria and Bacteroidetes increased, while Actinobacteria and Firmicutes decreased after the application of furfural residue. Pathogenic microbes *Alternaria, Trichocladium, Gibberella, Solicoccozyma, Bipolaris and Cladosporium* were not detected, and beneficial *Penicillium, Schizothecium* and *Rhizophlyctis* were detected in the soil after treatment with furfural residue.

The population of some pathogens decreased in the rhizosphere soil. Therefore, furfural residue could potentially be used for disease control. *MND1, RB41, UTCFX1, Nitrospira, Ellin6067, Adhaeribacter, Flavisolibacter, Polycyclovorans Haliangium*, and *Terrimonas* increased in the rhizosphere soil compared with the unplanted soil, while *Clostridium, Pseudarthrobacter* and *Roseiflexus* showed an opposite trend. The genera *Pirellula, Ellin6067* and *Luteimonas* decreased with plant growth. The application of furfural residues increased the abundance of microbes involved in nutrient absorption, organic matter decomposition and aromatic compound conversion. The composition of rhizosphere microorganisms shifted over time. Fertilization, particularly amendment with organic materials, is important to increase the shift in rhizosphere microbial communities in seed corn production system.

## Acknowledgement

We thank Yingmei Zhang at Lanzhou University for her suggestions for improving the manuscript. This work was financially supported by the Nature Science Foundation of China (41867010) and the program funds for the Longyuan innovative talents (2050205-1). We greatly thanks for Zhangye Zhongtian Agricultural science and technology Co. LTD for field plots.

