## Supplementary material for "Furfural residues as soil amendments for long-term seed corn production: effects on community composition and diversity of rhizosphere microbiota": Fig

A

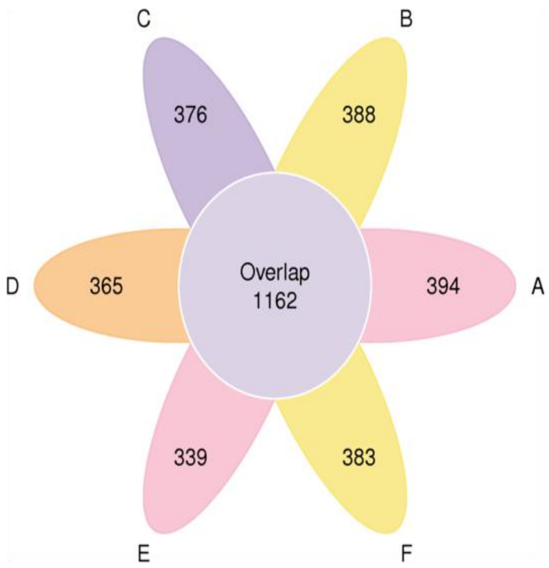


Fig. S1A Petal graph of bacteria OTUs presented in the soil of unplanted. The uppercase letter of A, B, C, D, E and F represents continuous cropping of corn seed production of 5, 10, 15, 20, 25 and 30 years, respectively.

B

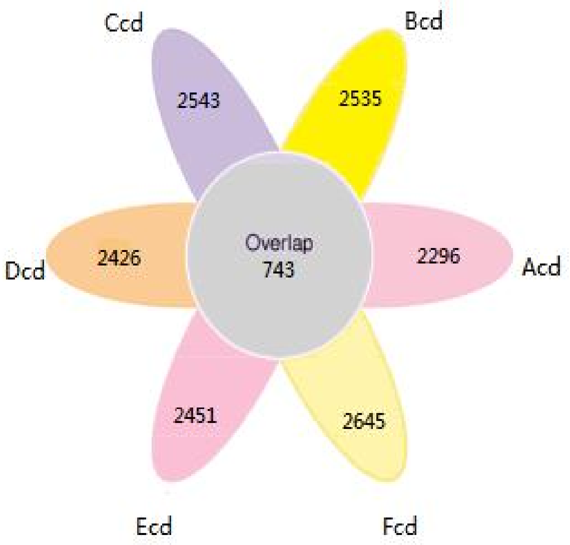


Fig.S1B Petal graph of bacteria OTUs presented in the soil of chemical fertilizer only. The lowercase letter cd means chemical fertilizer only at bell mouth stage. The uppercase letter of A, B, C, D, E, and F represents continuous cropping of corn seed production of 5, 10, 15, 20, 25 and 30 years, respectively.

C

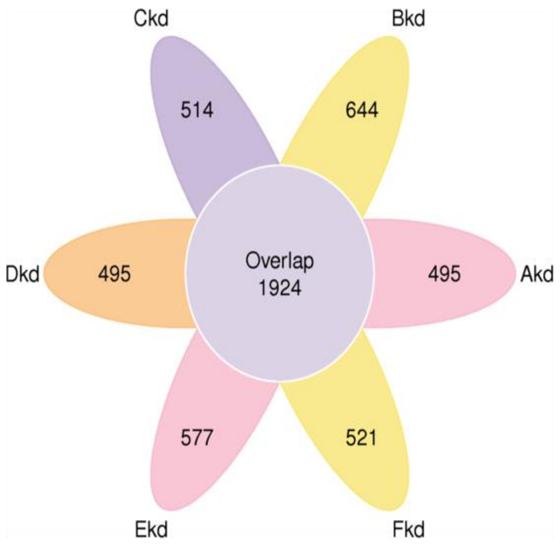


Fig.S1C Petal graph of bacteria OTUs presented in the soil of furfural residue application. The lowercase letter kd means furfural residue application at bell mouth stage. The uppercase letter of A, B, C, D, E and F represents continuous cropping of corn seed production of 5, 10, 15, 20, 25 and 30 years, respectively.

A

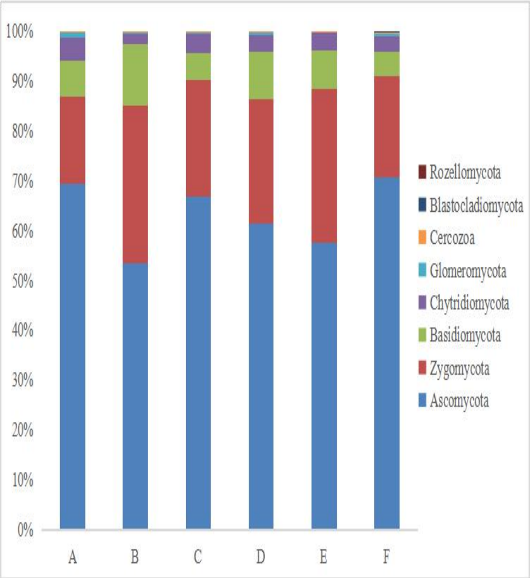


Fig.S2A Phylum of rhizosphere fungi of soil in March, which represents before planting. The uppercase letter of A, B, C, D, E and F represents continuous cropping of corn seed production of 5, 10, 15, 20, 25 and 30 years, respectively.

B

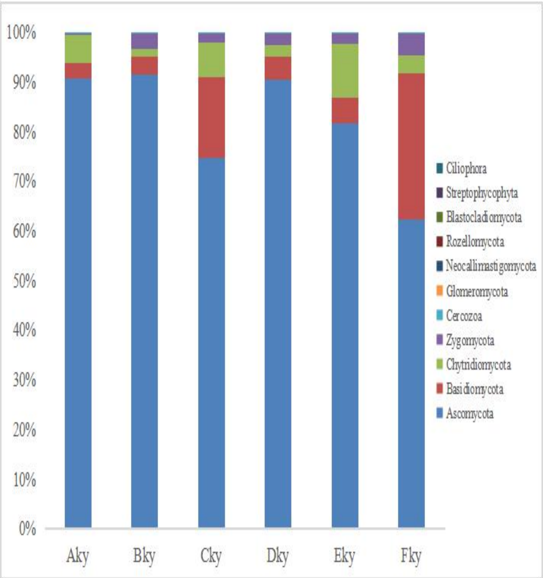


Fig.S2B Phylum of rhizosphere fungi of soil in May, which represents at seedling stage. The lowercase letter ky means furfural residue application at seedling stage. The uppercase letter of A, B, C, D, E and F represents continuous cropping of corn seed production of 5, 10, 15, 20, 25 and 30 years, respectively.

C

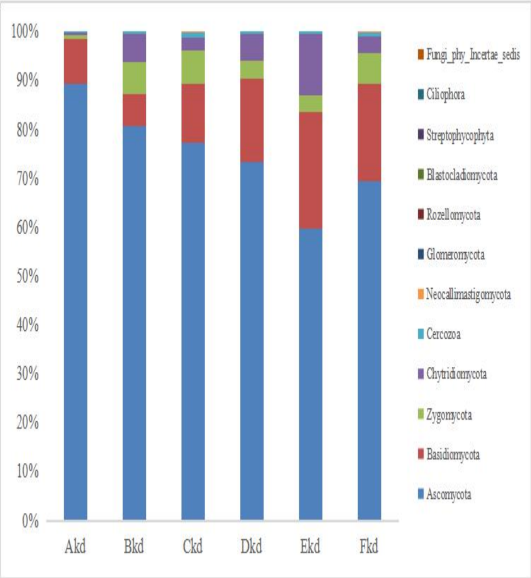


Fig.S2C Phylum of rhizosphere fungi of soil in March July, which represents at bell mouth stage. The lowercase letter kd means furfural residue application at bell mouth stage. The uppercase letter of A, B, C, D, E and F represents continuous cropping of corn seed production of 5, 10, 15, 20, 25 and 30 years, respectively.

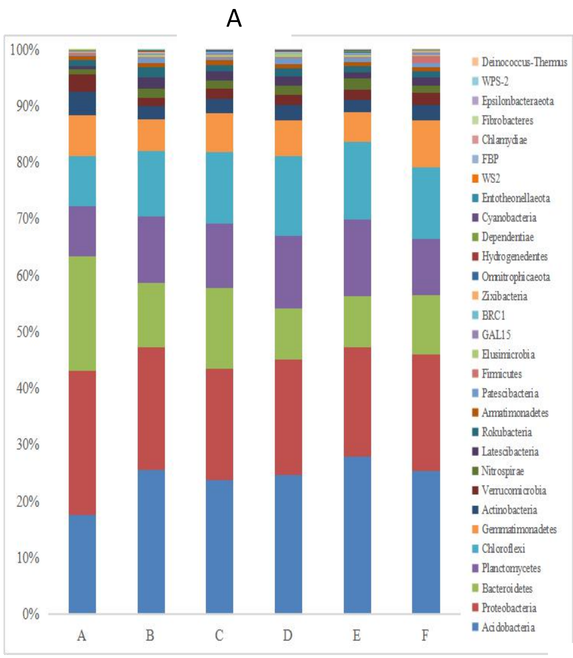


Fig.S3A Phylum of rhizosphere bacteria of soil in March, which represents before planting. The uppercase letter of A, B, C, D, E, and F represents continuous cropping of corn seed production of 5, 10, 15, 20, 25 and 30 years, respectively.

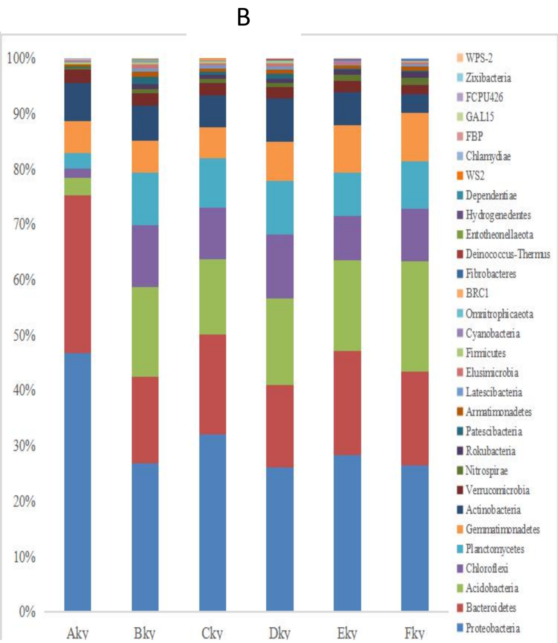


Fig.S3B Phylum of rhizosphere bacteria of soil in May, which represents at seedling stage. The lowercase letter ky means furfural residue application at seedling stage. The uppercase letter of A, B, C, D, E and F represents continuous cropping of corn seed production of 5, 10, 15, 20, 25 and 30 years, respectively.

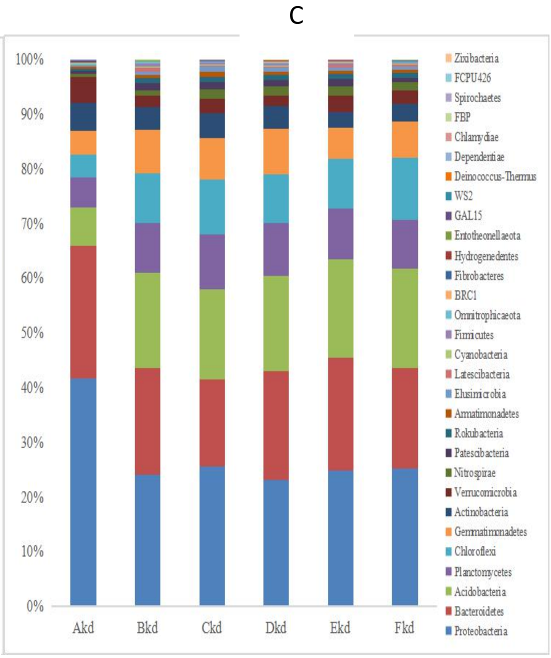


Fig. S3C Phylum of rhizosphere bacteria of soil in March July, which represents at bell mouth stage. The lowercase letter kd means furfural residue application at bell mouth stage. The uppercase letter of A, B, C, D, E and F represents continuous cropping of corn seed production of 5, 10, 15, 20, 25 and 30 years, respectively.
