## Supplementary material for "Furfural residues as soil amendments for long-term seed corn production: effects on community composition and diversity of rhizosphere microbiota": Table

Table S1 The properties of initial irrigated desert soil

| properties | OM  g. kg^-1^ | AN  g. kg^-1^ | AP  g. kg^-1^ | AK  g. kg^-1^ | pH |
| --- | --- | --- | --- | --- | --- |
| contents | 18.79~21.23 | 45.12~55.56 | 11.82~12.22 | 168.07~175.14 | 8.23~8.47 |

Note: OM means organic matter, AN means available-nitrogen, AP means available-phosphorus, AK means available-potassium.

Table S2 The geographic distribution of six study fields

|  | A | B | C | D | E | F |
| --- | --- | --- | --- | --- | --- | --- |
| Northern Latitude | 38˚49′33″ | 38°50′00″ | 38°49′54″ | 38°50′00″ | 38°49′54″ | 38°49′48″ |
| East Longitude | 100°20′01″ | 100°20′33″ | 100°20′33″ | 100°20′33″ | 100°20′32″ | 100°20′19″ |

Note: Uppercase letter of A, B, C, D, E and F means continuous cropping of 5, 10, 15, 20, 25 and 30 years, respectively.

Table S3 The characteristic of furfural residue after neutrilization

| Item | OM  % | HA  % | TN  % | TP  % | TK  % | pH |
| --- | --- | --- | --- | --- | --- | --- |
| contents | 76 | 11.5 | 0.6% | 0.08 | 1.18 | 7 |

Note: OM means organic matter, HA means humic acid, TN means total nitrogen, TP means total phosphorus, TK means total potassium, respectively.
